# Biomolecular condensates undergo a generic shear-mediated liquid-to-solid transition

**DOI:** 10.1101/2020.01.21.912857

**Authors:** Yi Shen, Francesco Simone Ruggeri, Daniele Vigolo, Ayaka Kamada, Seema Qamar, Aviad Levin, Christiane Iserman, Simon Alberti, Peter St George-Hyslop, Tuomas P. J. Knowles

## Abstract

A wide range of systems containing proteins have been shown to undergo liquid-liquid phase separation (LLPS) forming membraneless compartments, such as processing bodies^1^, germ granules^2^, stress granules^3^ and Cajal bodies^4^. The condensates resulting from this phase transition control essential cell functions, including mRNA regulation, cytoplasm structuring, cell signalling and embryogenesis^1–4^. RNA-binding Fused in Sarcoma (FUS) protein is one of the most studied systems in this context, due to its important role in neurodegenerative diseases^5–7^. It has recently been discovered that FUS condensates can undergo an irreversible phase transition which results in fibrous aggregate formation^6^. Gelation of protein condensates is generally associated with pathology. One case where liquid-to-solid transition (LST) of liquid-liquid phase separated proteins is functional, however, is that of silk spinning^8,9^, which is largely driven by shear, but it is not known what factors control the pathological gelation of functional condensates. Here we show that four proteins and one peptide system not related to silk, and with no function associated with fibre formation, have a strong propensity to undergo LST when exposed to even low levels of mechanical shear comparable to those found inside a living cell, once present in their liquid-liquid phase separated forms. Using microfluidics to control the application of mechanical shear, we generated fibres from single protein condensates and characterized their structures and material properties as a function of shear stress. Our results inform on the molecular grammar underlying protein LST and highlight generic backbone-backbone hydrogen bonding constraints as a determining factor in governing this transition. Taken together, these observations suggest that the shear plays an important role in the irreversible phase transition of liquid-liquid phase separated droplets, shed light on the role of physical factors in driving this transition in protein aggregation related diseases, and open a new route towards artificial shear responsive biomaterials.

---

During silk spinning in nature, the precursor protein molecules stored in liquid-liquid phase separated form undergo a LST to generate fibres mediated by mechanical shear^10–13^. It is increasingly recognised, however, that a wide range of other proteins can undergo LLPS inside the cell to form condensates^14,15^. This process is highly dynamic and crucial to cellular function^2,5,16^, while irreversible phase transitions lead to solid protein aggregates and can result in disease^6^. Yet, it remains unexplored whether functional protein condensates can also undergo a LST driven by mechanical shear.

In order to test the generality of fibre formation from liquid-liquid phase separated peptide and protein condensates, we selected a representative set of systems, including the RNA-binding proteins FUS and Ded1 from human and yeast cells, respectively, human membrane-binding protein Annexin A11, carboxybenzyl protected diphenylalanine (zFF) peptide^17^ and reconstituted silk protein. The monomeric solution of FUS protein is stable at high salt concentration (1 M KCl) and undergoes LLPS when the salt concentration drops below 150 mM^5^. We formed FUS droplets at protein and salt concentration of 2 µM and 50 mM respectively. By using a pair of tweezers, we were then able to pull solid fibres of 5-10 µm in diameter and 0.5-1 cm in length out from the droplet-containing solution (see Fig. 1). Next, we induced phase separation in a solution of Ded 1 protein by lowering the pH from 7.2 to 6.0 in PIPES buffer, whereas A11, zFF and silk protein were phase separated by mixing them with an aqueous solution of 10 % dextran and triggering fibre formation by applying shear as described above (Fig. 1a). Notably, FUS, Ded1, A11 and the dipeptide zFF, a short peptide composed of only two amino acids, exhibited shear induced fibre formation similar to that of the silk protein^8,9^, going through LLPS first and then LST, although they have no shared primary structure. However, fibres were not formed under shear with PR polypeptides (Fig. S1), which undergoes LLPS but exhibits no LST upon aging, due to lack of capability of backbone-backbone hydrogen bonding^18^.

**Figure 1:**
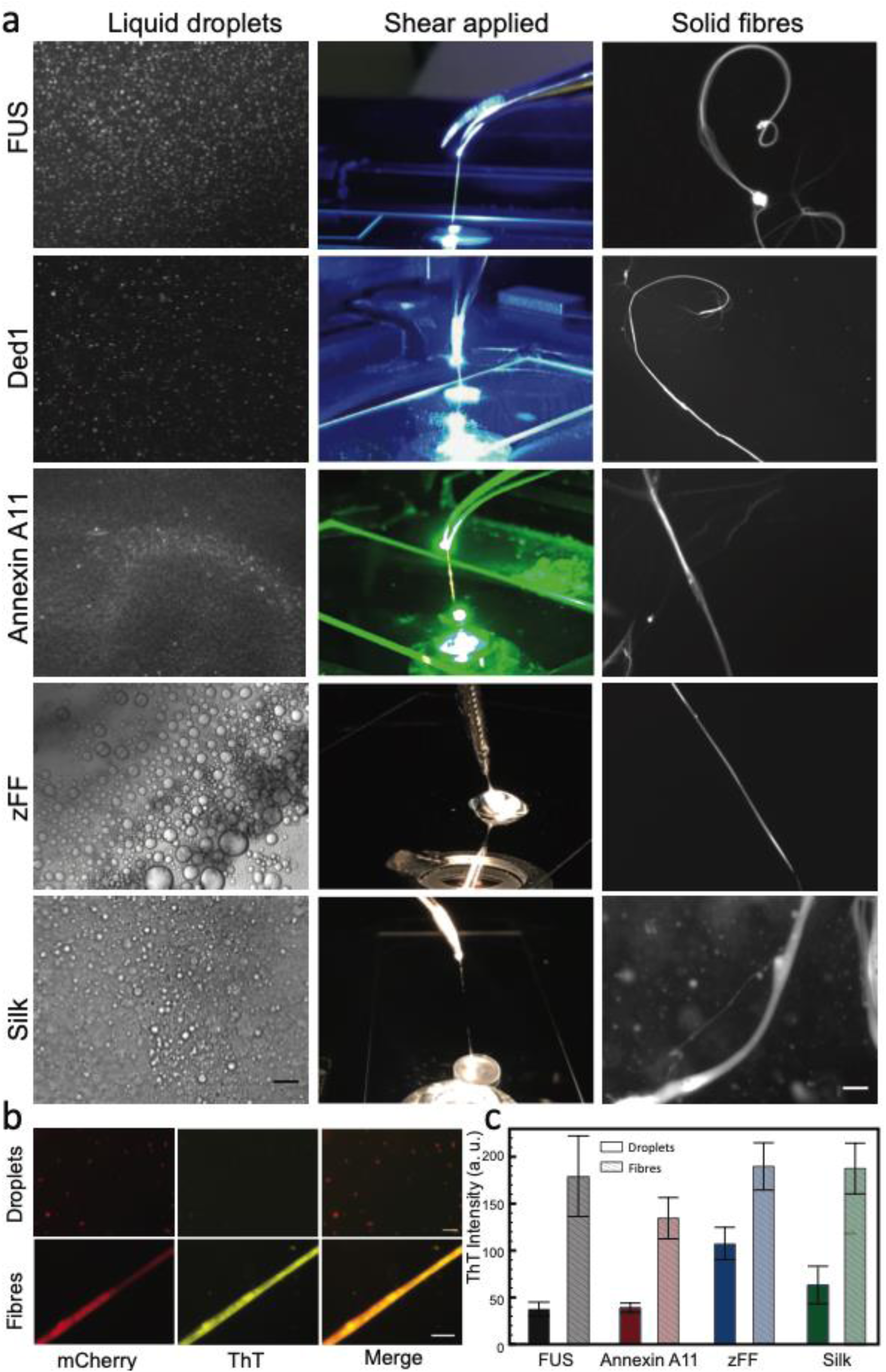
Generic fibre formation from liquid-liquid phase separated proteins and peptides. **a**. LLPS of FUS (GFP), Ded1 (GFP), Annexin A11 (mCherry), zFF and silk was induced by lowering the salt concentration, lowering the pH value and mixing with 10% dextran respectively (left column). The shear was applied by pulling the solution with a pair of tweezers (middle column). Fibre formation was observed by fluorescent microscopy (right column). zFF and silk fibre were stained with ThT. The scale bars of the first and third row of images are 40 μm. **b**. ThT staining of mCherry-tagged FUS droplets (top) and a fibre (bottom). Scale bars are both 5 μm. **c**. Fluorescence intensity of droplets and fibres after ThT staining for mCherry-tagged FUS (black), Annexin A11 (red), zFF (blue) and silk (green). n≥10 error bar SD. To be noted, as Ded1 is GFP-tagged, the excitation and emission wavelengths are overlapping with ThT staining, and thus not suitable for this assay.

We further studied the change in the secondary structure of the droplets prior to and following the application of shear. We first monitored qualitatively the structural changes of the droplets by staining mCherry-tagged FUS, Annexin A11 and unlabelled zFF and silk protein with Thioflavin T (ThT), which binds specifically to β-sheet rich structures. The FUS fibres obtained as a result, exhibited stronger ThT signal compared to that of liquid droplets immediately following LLPS (Fig. 1b), suggesting a structural transition from a disordered to a β-sheet rich ordered structure. The other proteins and peptide showed similar behaviour and the difference in ThT intensity of droplets and fibres is quantified and summarized in figure 1c. Next, we performed conventional Fourier Transform Infrared spectroscopy (FTIR) measurements and analysis on bulk samples before and after the application of shear to further systematically investigate the change in protein structure. Following shearing of the bulk solution, we observed a significant shift of the Amide band I to lower wavenumbers for FUS, Ded1 and silk protein, suggesting the formation of hydrogen bonding and structural rearrangement (Fig. S2). However, due to the heterogeneity of the bulk samples (mixture of fibres and droplets), the change was not significant for Annexin A11 and zFF.

In order to exclude the influence from sample heterogeneity and investigate the structural transitions occurring during fibre formation at a single droplet and single fibre resolution, we focused on the FUS protein system and exploited the capabilities of polarized infrared nanospectroscopy (AFM-IR). The AFM-IR method combines the high-spatial resolution of scanning probe microscopy (AFM ∼ 10 nm) with the chemical and structural recognition power of IR spectroscopy (Fig. 2a). By using AFM-IR, we acquired the maps of 3D morphology, IR absorption and stiffness of the FUS samples with nanoscale resolution^7,20,21^. We first acquired a morphology map of a single droplet (Fig. 2b). We then simultaneously acquired the morphology (Fig. 2c), IR absorption in the Amide I band at 1655 cm^-1^ (Fig. 2d) and nanomechanics (Fig. 2e) of a single fibre. The comparison of the average spectra on a droplet and on a fibre formed through the application of shear shows that the Amide band I and II of the fibre are shifted toward lower wavenumbers than in the droplet, indicating the formation of intermolecular hydrogen bonding (Fig. 2f). Moreover, the appearance of a shoulder in the Amide I band at 1625 cm^-1^ indicates the formation of intermolecular β-sheet, in good agreement with the FTIR results from the bulk samples (Fig. S2a). The relative change in the secondary structure of droplets and fibres is presented in figure 2g. These results show that there was more relative formation of parallel β-sheet and α-helix content, and less antiparallel β-sheet and random coil in the fibres.

**Figure 2:**
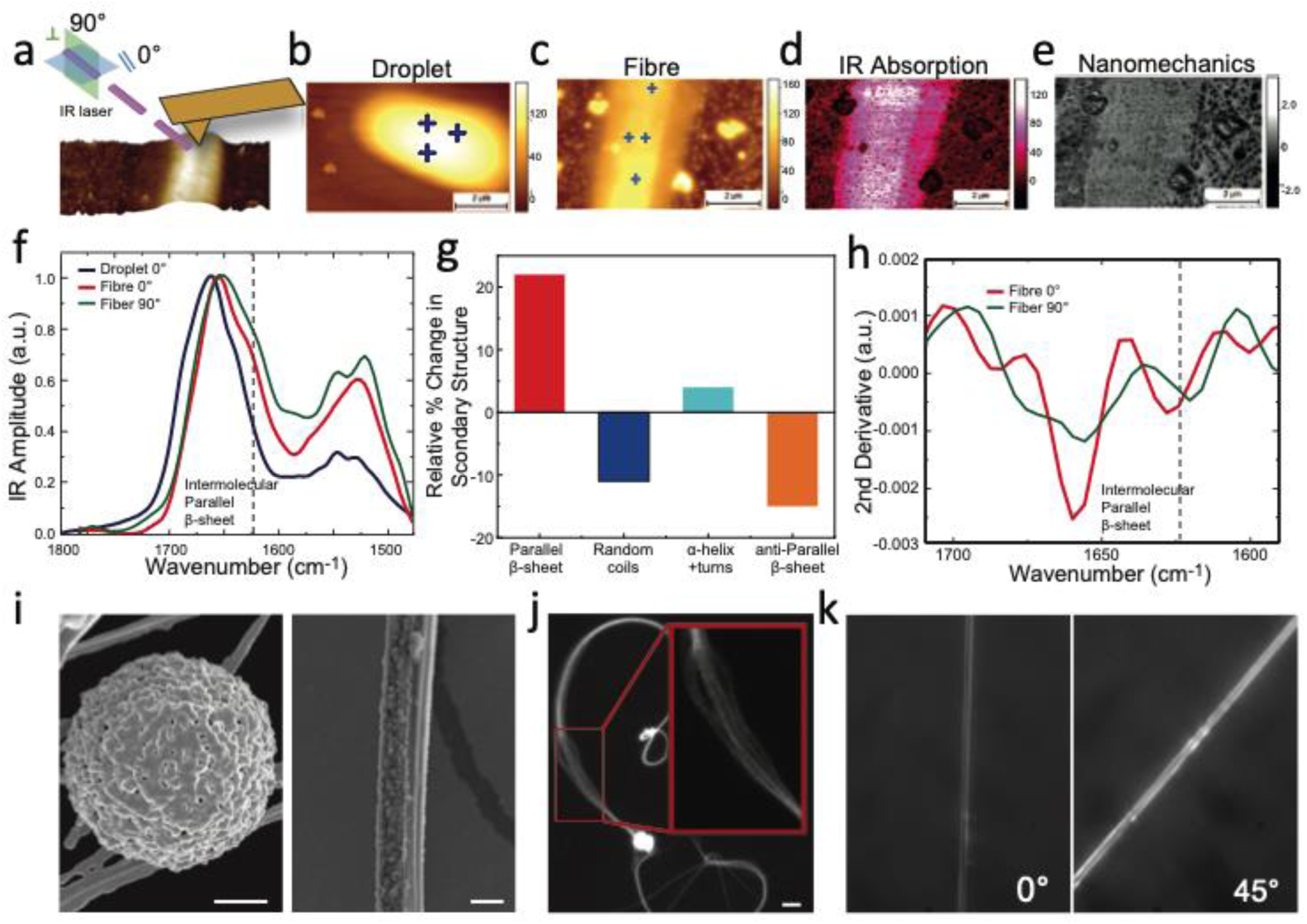
Structural changes in FUS protein droplets following shear. **a.** Scheme of the measurement setup for polarized infrared nanospectroscopy (AFM-IR) on a single fibre with polarized light source parallel (0°) and perpendicular (90°) to the deposition surface. 3D morphology map of a droplet (**b.**) and fibre (**c.**) Crosses indicate the locations of the spectroscopy measurements. **d.** Map of IR absorption in the Amide I band (1655 cm^-1^). **e.** Maps of the nanomechanical properties of the fibre^7,19^. **f**. Nanoscale localized IR spectrum as a function of the polarization angle of a droplet at 0° (blue line), fibre at 0°(red line) and 90° (green line). Spectra were acquired at the positions indicated by the blue crosses. **g.** Relative change in secondary structure between droplets and fibres **h**. Second derivative of the IR spectrum of the fibre with polarization 0° (red line) and 90° (green line). **i.** Scanning electron microscopy (SEM) images of a droplet aged for 1h (Scale bar 500 nm) and a fibre (scale bar 2 μm). **j**. Fluorescence microscopy image of a FUS fibre in 50 mM tris buffer. Aligned nanofibrils within the fibre are shown in the zoomed in insert. The scale bar is 20 μm. **k.** Polarization microscopy images of a fibre at 0° and 45°.

To obtain further structural information on the fibrillar state, we compared the AFM-IR spectra acquired with polarized light source parallel (0°) and perpendicular (90°) to the deposition surface. The IR spectrum of the fibre changed as a function of the polarization. In particular, with an angle of 90°, a stronger shift at lower wavenumbers of the Amide band I and a more pronounced shoulder at 1625 cm^-1^ were detected (Fig. 2f). This change was more pronounced once we performed a second derivative analysis of the IR spectra at 0° and 90°. The structural contributions changed in intensity as a function of the polarized light, in particular, with a relative increase in the intermolecular parallel β-sheet content (1625 cm^-1^) at 90° (Fig. 2h). This polarization effect demonstrates that the structure of the fibre is anisotropically ordered and that there are more intermolecular β-sheets oriented perpendicular to the fibre axis^22^. We next performed scanning electron microscopy (SEM) to characterize the structural morphology of the FUS phases. The micrographs revealed a porous network for FUS droplets and elongated rod-shaped fibres with aligned nanofibrils (Fig. 2i). The alignment of the nanofibrils within the fibre was also observed through fluorescence microscopy (Fig. 2j). This anisotropy was further confirmed by polarization microscopy, which showed a maximum birefringence signal at 45°, indicating that the fibres contain highly aligned nanofibrils at their core (Fig. 2k).

In order to explore this phenomenon in a quantitative manner, a microfluidic platform was developed to induce phase separation, apply precise shear stress and characterize the mechanical properties of the formed structures in a liquid environment. In particular, through this approach, we induce and control the phase separation in the channel so that the droplets remain in a liquid state prior to the application of shear. The design of the microfluidic device used is illustrated in Figure 3a. Two large flow chambers were connected by 5 series of small bridge channels with dimensions of 3, 5, 7, 11 µm in width and 4.2 µm in height (Fig. 3a). By modulating the flow rates in the two large chambers, different pressure drops were obtained across the small bridges to generate variable levels of shear. We first induced phase separation by flowing FUS protein in a high salt buffer in the right chamber and low ionic strength buffer in the left chamber. The flow rates were chosen to generate a small imbalance in the pressure between left and right chambers to drive the protein solution through the bridges and allow phase separation to be induced in the low salt environment at the entrance of the bridge channels (Fig. 3b). Time lapse microscopy of the process revealed the nucleation and growth of FUS droplets. Within 2 minutes, the droplets grew to i.e. 3 µm in diameter (Fig. 3b). We then rapidly applied a higher flow rate to increase the level of shear. By increasing the average shear stress from 0.06 to 0.5 Pa across the small bridges, a liquid droplet could be seen to deform and elongate along the bridge channel (Fig. 3c left). The pressure difference was then switched off by applying the same flow rates in both large chambers. Following this, the formed fibre was first seen to retract and then curl back inside the bridge, indicating the formation of an elastic fibre from a FUS protein droplet (Fig. 3c right). We obtained the detailed flow field when the fibre was formed and retracted by performing 3D numerical simulation as shown in Figure 3c top. We further quantified the effect of the shear stress on the phase transition of FUS droplets by applying different flow rates to form the fibres within the microfluidic channel. Detailed flow fields at each average shear stress is shown in Figure S3 and we found a critical shear stress 0.5 Pa was needed to trigger the fibre formation from droplets.

**Figure 3.**
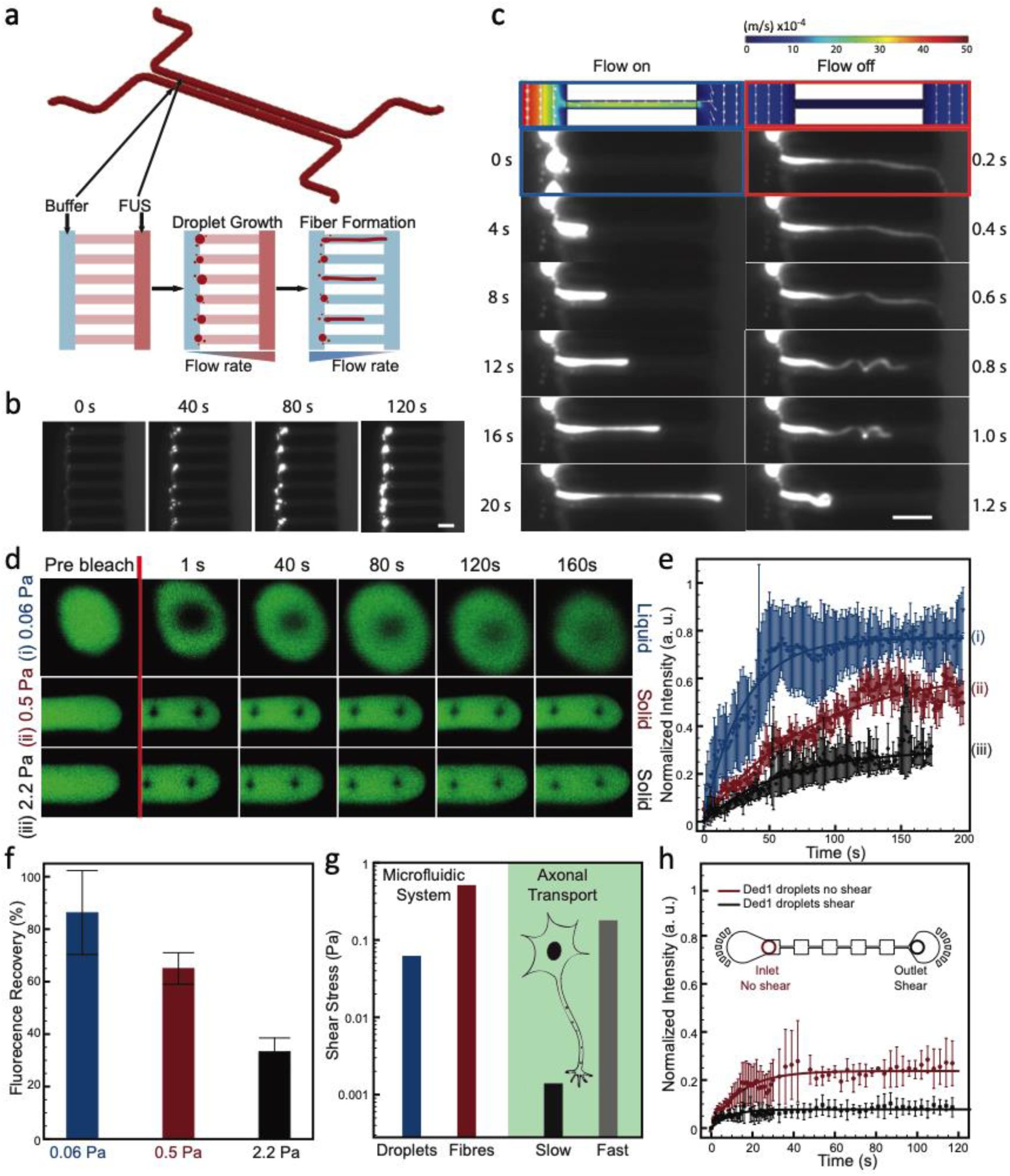
Shear mediated fibre formation probed by microfluidics. **a.** The design of the microfluidic device and the schematic of the droplets’ growth and fibre formation under varied flow conditions. Two large flow chambers (with width × height, 50 × 25 µm) were connected with 5 series of small bridges (with 3 × 4.2, 5 × 4.2, 7 × 4.2, 9 × 4.2, 11 × 4.2 µm width × height) **b.** Time dependant fluorescence images of FUS droplets formation at the entrance of the small bridge channels, scale bar 5 µm. **c.** A single fibre was formed from a single droplet under shear with the flow on, which then retracted and curled when the applied pressure was turned off (Mov.1), scale bar 5 µm. 3D numerical simulation of the flow field when the shear was applied (top row). **d.** Images **e.** Curves and **f.** Average fluorescence recovery of FUS droplets and fibres formed at different shear stress: (i) 0.06 Pa, (ii) 0.5 Pa and (iii) 2.2 Pa, n≥3 error bar SD **g**. Comparison of the shear stress present in the human axonal transport^23–26^ relative to the regime explored in this study. **h.** Fluorescence recovery after bleaching of Ded1 droplets before and after having experienced the shear in the constricted microfluidic channel, n≥3 error bar SD.

We further characterized the rheology of the droplets and fibres by Fluorescence Recovery After Photobleaching (FRAP) measurements, which were used to determine the diffusion coefficient of the molecules in the droplets and hence report on the local viscosity^27^. We found that the FUS droplets remain liquid and the fluorescence recovery reached up to ∼85% when a shear stress upto 0.06 Pa was applied. Once these droplets were deformed into solid fibres by applying a critical shear stress of 0.5 Pa, only ∼65% of the fluorescence was found to be recovered. Finally, by applying an even higher shear stress of 2.2 Pa, the formed fibres only recovered ∼35% of their initial fluorescence intensity (Fig. 3d–f). It is interesting to note that these shear stress values are comparable to the biologically relevant range. In fact, the shear stress that can be generated during human axonal transport ranges from 0.001 to 0.18 Pa^23–26^ (considering the diameter of axons of about 1 μm and viscosity ∼5 cP^28^) (Fig. 3g), highlighting the fact that biological systems operate close to the critical shear value that the present work has revealed can trigger a transition to an irreversible solid phase associated with malfunction. These results demonstrate that a critical shear stress is required to trigger the fibre formation. Additionally, the higher the shear stress applied, the more compact the structure of the fibre obtained, showing the significance of shear on protein condensates phase transition. As another example, Ded 1 was tested under shear by flowing it through a channel with series of expansions and constrictions right after LLPS (Fig. 3h). The droplets at the inlet of the channel not exposed to shear had higher fluorescence recovery (∼22%) than the ones at the outlet that had been exposed to shear (∼7%) after bleaching (Fig. 3h), showing a similar shear response as FUS protein. Furthermore, we characterized in more detail the tensile strength of a tweezer-pulled FUS fibre by applying an increasing load until the point of fibre fracture (Fig. 4a). The results showed a maximum tensile strength of 35 MPa, a toughness of 2.8 MJ/m^3^ and a Young’s modulus of approximately 3 GPa, which is comparable to silk fibres from cocoons^29,30^ (Fig. 4b). This similarity can be rationalized by the fact that both fibrous materials are hold together with a dense intermolecular β-sheet network. Moreover the very high level of mechanical stiffness and strength highlights the potentially damaging effect of these structures once formed in the intercellular milieu.

**Figure 4:**
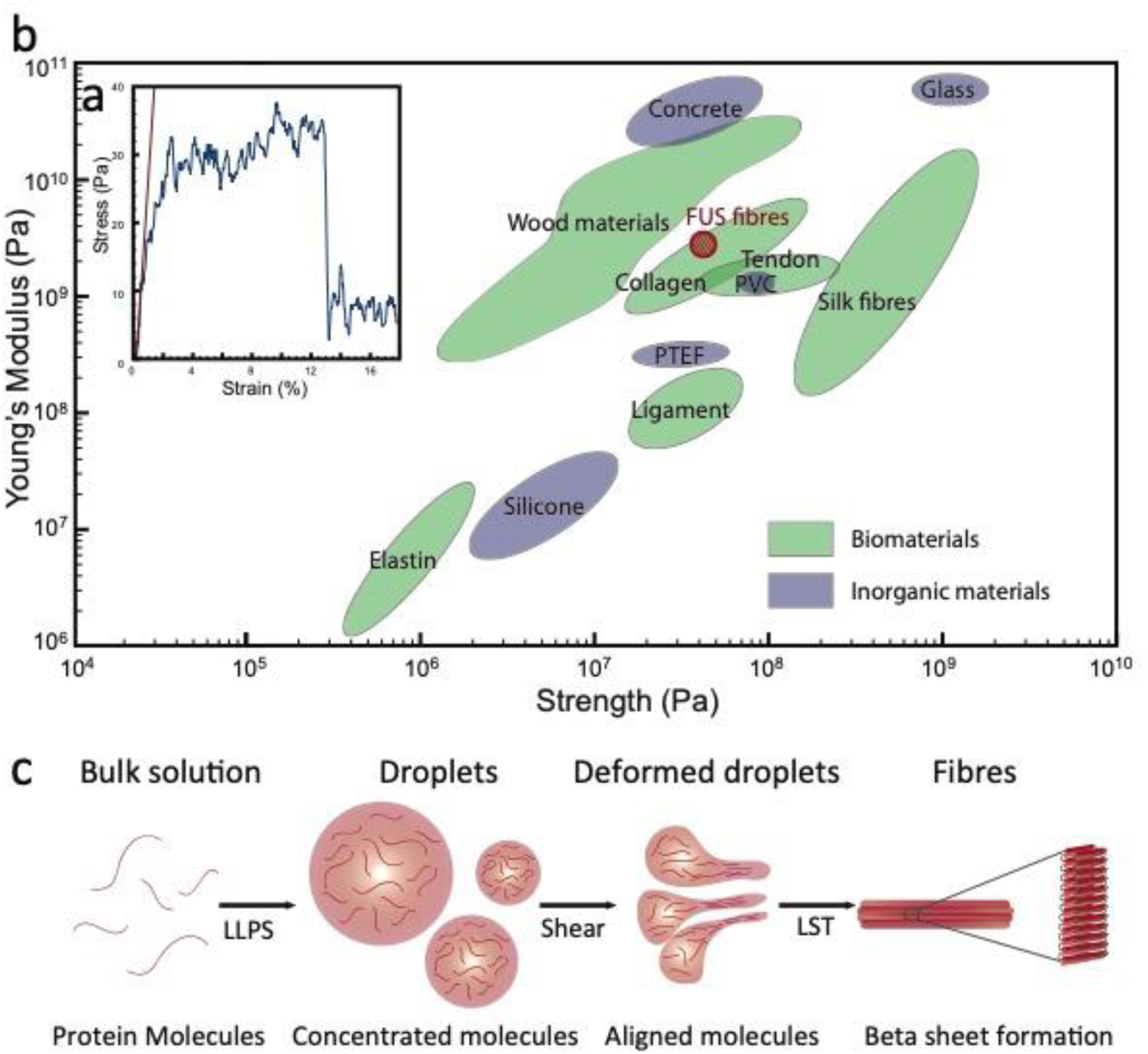
Material properties of fibres and proposed model of fibre formation from protein condensates under shear. **a.** Tensile strength measurement of a tweezer-pulled dried FUS fibre. **b.** Young’s modulus versus strength of the FUS fibre compared to a range of biological and inorganic materials. **c.** Protein molecules get concentrated in the droplets after LLPS. Shear aligns the molecules/oligomers inside the droplets, decreases their excluded volume, promotes inter-molecular interactions resulting in the formation of fibres rich in β sheets.

Proteins, once in condensate form, get concentrated and have a higher viscosity than that of dilute protein solution^6^, implying that protein molecules within droplets experience greater mechanical stress at an identical shear rate. The application of shear stress can thus align the protein molecules, promote inter-molecule interactions and trigger a phase transition that results in the formation of microscale fibres composed of aligned nanofibrils rich in intermolecular hydrogen bonded β-sheets (Fig. 4c).

To conclude, we report here a general shear response of peptides and proteins that can form solid fibres from phase separated liquid protein droplets. This process has striking similarities with that of silk fibre formation, yet is independent of the protein/peptide primary sequence. This phenomenon was probed with an array of microfluidic based approaches to allow for controlled LLPS and tuneable shear stress to be applied, inducing fibre formation from single droplets while minimizing the influence of any air-water interfaces. With this methodology we were able to achieve shear conditions comparable to those in cells. This observation suggests that biology has most likely evolved mechanisms to protect against shear-mediated LST, and it is interesting to speculate that the heterogeneity nature of many naturally occurring condensates can contribute to this protective effect. Our findings unveil the importance of mechanical shear for protein phase transitions that drive liquid condensates to form solid fibres, shedding light on the possible mechanisms of disease-related protein aggregation.

## Methods

### Proteins/Peptide sample preparation

#### FUS

cDNA encoding FUS (amino acid residues 1-526) was cloned into a baculovirus expression system vector, pACEBac2, with a protease cleavable N-terminal MBP tag and a C-terminal GFP-6xHis tag. The purified recombinant bacmid encoding FUS was transfected into Sf9 insect cells. The protein was expressed for 4 days post infection with the baculovirus harbouring the recombinant bacmid. The cell culture was harvested by centrifugation at 4,000 rpm for 30 min. The cell pellet was homogenised into the lysis buffer containing 50 mM Tris, 1 M KCl, 0.1% CHAPS, 1 mM DTT, 5% glycerol at pH 7.4. Post lysis the sample containing the protein was spun at 40,000 rpm in an ultracentrifuge to remove the cell debris. The supernatant was collected and processed using three steps purification protocol including, Ni-NTA affinity column, amylose affinity column and the final polishing step of size exclusion chromatography in the buffer containing 50 mM Tris, 1 M KCl, 1 mM DTT, 5% glycerol, pH 7.4. The final purity of the protein was greater than 95%. LLPS was induced by diluting the FUS solution in the 50 mM Tris buffer at final concentration 2 µM (FUS) and 50 mM (KCl) respectively.

#### Annexin A11

cDNA encoding the full length Annexin A11(amino acid residues 1-505) was cloned into pACEBac2 vector with a TEV cleavable N-terminal MBP tag and a C-terminal Cherry-His tag. The protein was expressed and purified from insect Sf9 cells. After 6 days of infection, cells were harvested and lysed by homogenising into a resuspension buffer containing 50 mM HEPES, 100 mM NaCl, 1 mM EDTA, 5% glycerol, 0.1% CHAPS, pH 7.4. Protein purification was undertaken as described above for FUS protein, using a three-step purification scheme. The final buffer used for Annexin A11 was 50 mM HEPES, 225 mM NaCl, pH 7.4. The final purity of the protein was greater than 95%. An Annexin A11 solution at concentration of 10 μM was mixed with dextran to induce the LLPS.

#### DED1

cDNA encoding Ded1 was cloned into a baculovirus expression system vector, pOCC120, with a GST protease cleavable N-terminal MBP tag and a C-terminal GFP and GST protease cleavable 6xHis tag. SF9 insect cells were transfected with recombinant baculovirus stocks and incubated for four days. The cells were harvested by centrifugation (15 min 500 x g), resuspended in 30 mL of lysis buffer (50 mM Tris/HCl, pH 7.6, 1 M KCl, 1 mM DTT, 1 protease inhibitor tablet with EDTA/50 ml buffer, 2 mM EDTA, 10 µl benzonase (250U/mL) and lysed using the EmulsiFlex-C5 (Avestin, Ottawa, Canada). The lysates were clarified by centrifugation (16,000 rpm for 1 h at 4°C, rotor JA 25.50, Beckman Coulter, Brea, California, USA). The supernatant was incubated with amylose resin for 1 h at 4°C and loaded onto a 20 ml chromatography column (Bio-Rad, Hercules, CA, USA). After washing 3 column volumes with wash buffer (50 mM Tris, pH 7.6, 1 M KCl, 1 mM DTT, 2 mM EDTA), the protein was eluted with elution buffer (wash buffer plus 20 mM maltose). Dialysis of the protein as well as cleavage of the tags (His and MBP) was performed overnight at 4°C in wash buffer with 0.01 mg/ml GST-3C Protease. The protein aggregates were removed by centrifugation in Falcon Tubes for 4 min at 3452 x g. The protein was concentrated using Amicon Ultra-15-30k centrifugal filters at 3452 x g. The proteins were loaded onto an Äkta Pure chromatography setup (GE Healthcare, Uppsala, Sweden) equipped with a Superdex 200 26/60 column or Superdex 200 16/60 column (GE Healthcare, Piscataway, NJ, United States). After SEC, the protein was concentrated by centrifugation using Amicon Ultra 15-30k centrifugal filters (Merck, Kenilworth, NJ, USA) at 3452 x g. The protein was flash-frozen using liquid-nitrogen and stored at −80°C.

#### zFF

Carboxybenzyl (Z)-protected diphenylalanine (zFF) dipeptide was purchased from Bachem (Switzerland). Z-FF was dissolved to a concentration of 100 mg/ml in DMSO and was then diluted further into a 10 % dextran solution to induce the LLPS.

#### PR25

The peptide with 25 repeats of proline-arginine (PR25) was purchased from GenScript (Hong Kong). PR25 was dissolved in water and then mixed with polyU at final concentration 100 μM and 1 μg/μl respectively to induce the LLPS.

#### Reconstituted Silk Fibroin (RSF) preparation

Silk protein was prepared based on a well-developed protocol^31^. Bombyx mori silk cocoons (Mindsets (UK) Limited) were cut into pieces and placed in a beaker containing a solution of 0.02 M sodium carbonate. The solution was boiled for 30 minutes and then the insoluble fibroin was removed from beaker. The fibroin was rinsed with water for three times and left for 3 days at room temperature to dry out. A 9.3 M lithium bromide solution was added to the dried silk fibroin in a 1:4 ratio of silk fibroin to lithium bromide. The mixture was heated at 65 °C for 4 h. The resulted solution was then dialysed against Mili-Q water in order to remove LiBr. The solution was left for 48 h at 4 C° while changing the water for 6 times in total. The dialyzed solution was then centrifuged at 9,000 rpm at 4 C° for 20 min to remove the remaining impurities. The centrifuge process was repeated twice, and the final solution was stored at 4 C°. A silk solution at concentration of 110 μg/ml was mixed with dextran to induce the LLPS.

### Thioflavin T (ThT) staining

The β-sheets formation in the droplets and fibres was monitored by ThT staining. A final concentration of 100 μM ThT was used. The fluorescence images were taken under a microscope using a filter with excitation wavelength of 440 nm and an emission wavelength of 480 nm.

### Scanning Electron Microscopy (SEM)

A solution of liquid-liquid phase separated FUS protein was aged and sampled at 1 min, 1 h and 5 h. The samples were then briefly dipped twice in de-ionised water to remove any buffer salts and quickly plunge-frozen by dipping into liquid nitrogen-cooled ethane. Then, samples were freeze-dried overnight in a liquid nitrogen-cooled turbo freeze-drier (Quorum K775X). After that, the samples were mounted on aluminium SEM stubs using conductive carbon sticky pads (Agar Scientific) and coated with 15 nm iridium using a Quorum K575X sputter coater. The images were taken by using a FEI Verios 460 scanning electron microscope run at 2 keV. Secondary electron images were acquired using a high resolution Through-Lens detector in full immersion mode. Fibres were pulled from the LLPS solution and directly air dried and coated with 10 nm platinum. MIRA 3 FEG-SEM (TESCAN) was used in the 2 kV SE mode to take the images.

### Flow field simulation

Numerical simulations were performed using COMSOL Multiphysics 5.2a (Massachusetts, USA). To obtain the 3D flow field of the complete microfluidic device the one-phase laminar flow physics was used. We modelled the aqueous protein suspension as water (via the built-in material’s library) and used no-slip boundary conditions at the walls. The flow rate at each inlet was set as a laminar inflow and the outlets were set at zero pressure with suppressed backflow. The velocity and pressure were extracted from a surface parallel to the xy plane at z = 2 µm, correspondent to the middle plane of the narrow bridges. The mean shear rate was obtained by averaging the shear rate values extracted from the cross-section at the centre of a narrow bridge. The shear stress is then simply obtained from the shear stress by multiplying it by the dynamic viscosity.

### FTIR

Attenuated total reflection infrared spectroscopy (ATR-FTIR) was performed using a Bruker Vertex 70 spectrometer equipped with a diamond ATR element. The resolution was 4 cm^-1^ and all spectra were processed using Origin Pro software. The spectra were averaged (3 spectra with 256 co-averages), smoothed applying a Savitzky-Golay filter (2^nd^ order, 9 points) and then the second derivative was calculated applying a Savitzky-Golay filter (2^nd^ order, 11 points).

### Infrared Nanospectroscopy

A nanoIR2 platform (Anasys, USA), which combines high resolution and low noise AFM with a tunable quantum cascade laser (QCL) with top illumination configuration was used. The samples morphology was scanned by the nanoIR microscopy system, with a rate line within 0.1-0.4 Hz and in contact mode. A silicon gold coated PR-EX-nIR2 (Anasys, USA) cantilever with a nominal radius of 30 nm and an elastic constant of about 0.2 N m^-1^ was used. To study polarisation effects, the IR light was polarized parallel and perpendicular the surface of deposition^22^. All images were acquired with a resolution of at least 500×500 pixels per line. The AFM images were treated and analysed using SPIP software. The height images were first order flattened, while IR and stiffness related maps where only flattened by a zero-order algorithm (offset). Nanoscale localised spectra were collected by placing the AFM tip on the top of the FUS droplets or fibres with a laser wavelength sampling of 2 cm^-1^ with a spectral resolution of 4 cm^-1^ and 256 co-averages, within the range 1400-1800 cm^-1^ ^32^. Within a droplet or a fibre, spectra where acquired at a least 3 different nanoscale localised positions, the spectrum at each position being the average of 5 spectra. The average spectrum of the different was subtracted by the baseline signal of the substrate and salt^33^. Successively, the spectra were smoothed by Savitzky-Golay filter (second order, 9 points) and normalized. Spectra second derivatives were calculated and smoothed by Savitzky-Golay filter (second order, 9 points). Spectra were analysed using the microscope’s built-in Analysis Studio (Anasys) and OriginPRO. All measurements were performed at room temperature and with laser power between 1-4% of the maximal one and under controlled Nitrogen atmosphere with residual real humidity below 5%.

### Microfluidics

The microfluidic channels were fabricated based on previous protocols, using polydimethylsiloxane (PDMS) (Sylgard 184 kit, Dow Corning, Midland, MI, USA). The channels were plasma-treated and prefilled with buffer. The samples were flown in with precisely controlled flowrate with a syringe pump (Cetoni neMESYS, Cetoni GmbH, Korbussen, Germany). The bright field and fluorescence images were obtained from an inverted microscope (Axio Oberver A1, Zeiss, Cambridge, UK).

### Fluorescence Recovery After Photobleaching (FRAP)

FRAP was performed on a confocal microscope (Leica TCS SP8, Leica Microsystems GmbH, Wetzlar, Germany) by using spot bleaching mode. The fluorescence intensity was detected over time after bleaching and normalized with the signal from the whole sample.

### Tensile strength analysis

Tensile testing was performed using Tinius Olsen 5 ST with 5 N load cell, with the crosshead speed of 1 mm/min. In order to calculate stress, the obtained force data was divided by the cross-section area of the fibre where the diameter was measured prior to the testing using optical microscopy. The stress data was smoothed taking moving average of 24 neighbouring elements (corresponding to 0.14 % of strain) in order to reduce noise.

## Acknowledgements

This work is supported by the Welcome Trust, ERC, Alzheimer Association Zenith, ALS Canada-Brain Canada, Canadian Institutes of Health Research and the Cambridge Centre for Misfolding Diseases. The authors would like to thank S. Zhang, Y. Lu and K. L. Saar for their help in fabrication of the microfluidic devices. K.H. Muller for the help of flash freezing and SEM imaging.

## Authors Contributions

Y.S. and T.P.J.K. conceived and designed the study. Y.S., S.F.R., A.K. and A.L. performed the experiments. Y.S., S.Q., P.StGH., C.I., S.A. and A.K. produced the materials. S.F.R. performed AMF-IR and analyzed the data. D.V. simulated the flow field. Y.S. imaged the samples under SEM, fluorescent microscope, ran microfluidic experiments and FRAP analysis. A.K. performed tensile strength measurement. Y.S., S.F.R., D.V., A.K. P.StGH, S.A. and T.P.J.K. analysed the data. All the authors contributed to the writing of the manuscript.

## Competing financial interests

The authors declare no competing financial interests.

## Notes

#### Summary of Updates

Minor corrections of text in the conclusion section.

