## Supplementary information for "Biomolecular condensates undergo a generic shear-mediated liquid-to-solid transition"

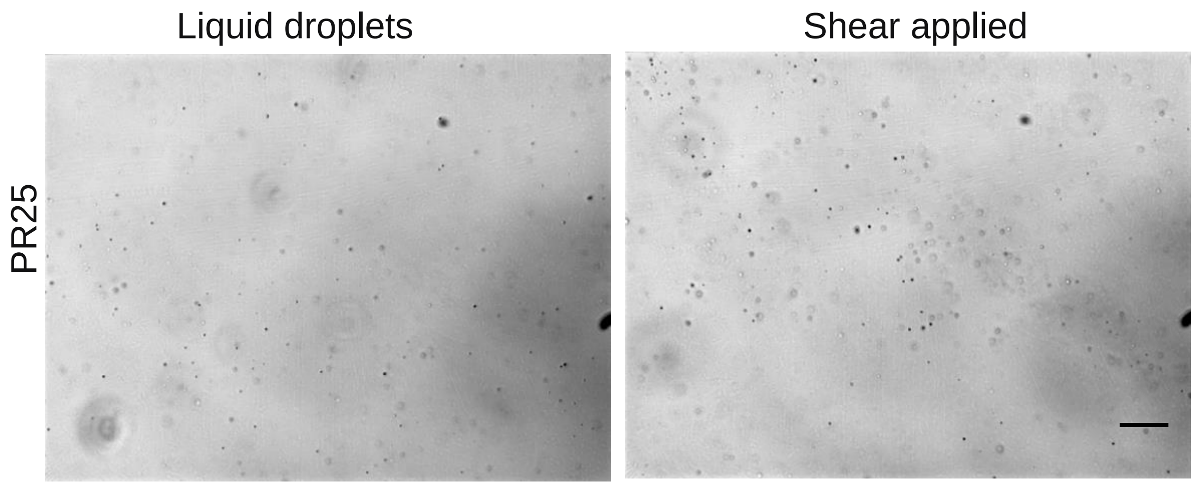


Figure S1: LLPS of PR25 before and after shear, scale bar is 40 μm.

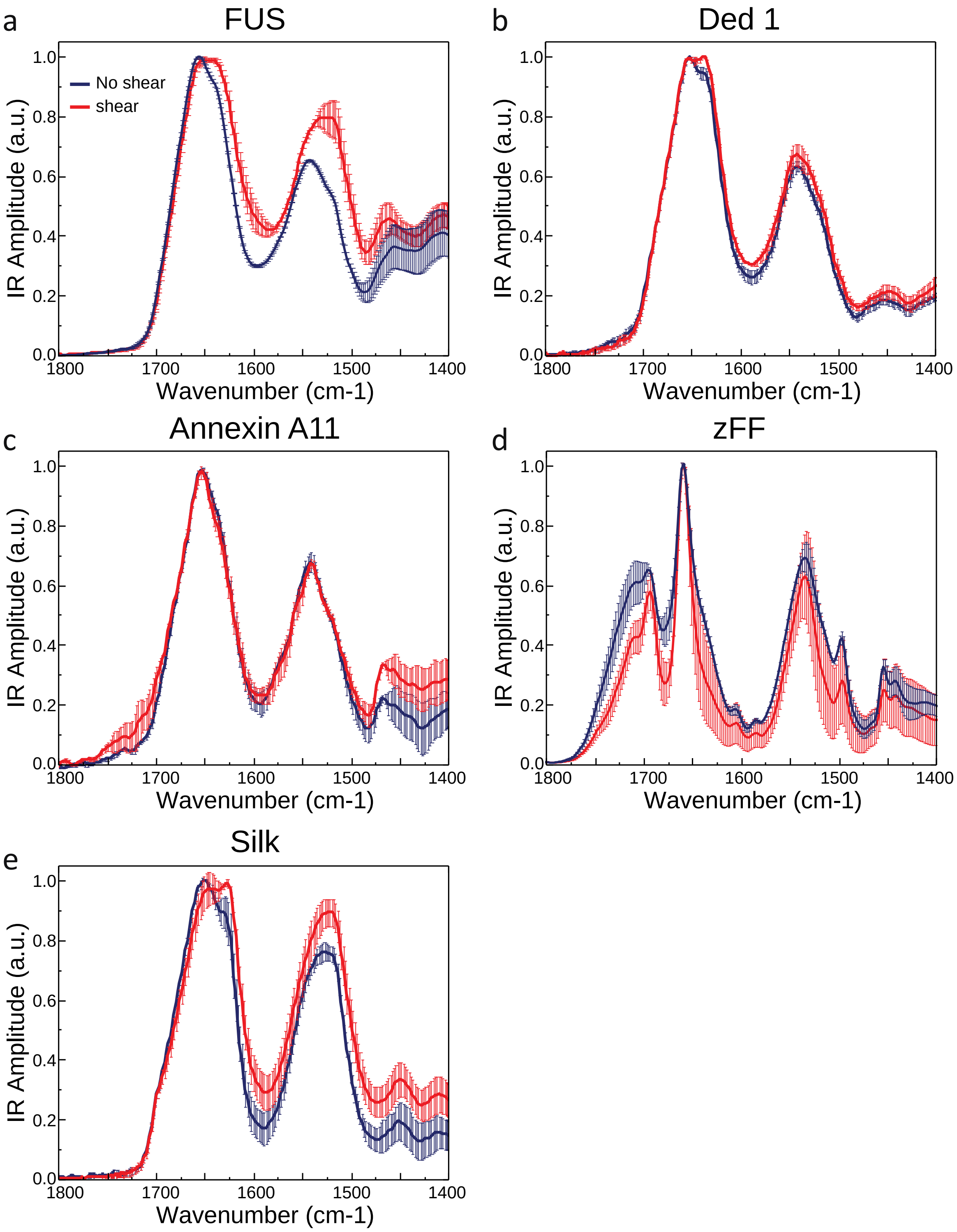


Figure S2: IR spectrum of liquid-liquid phase separated FUS solution before and after shear.

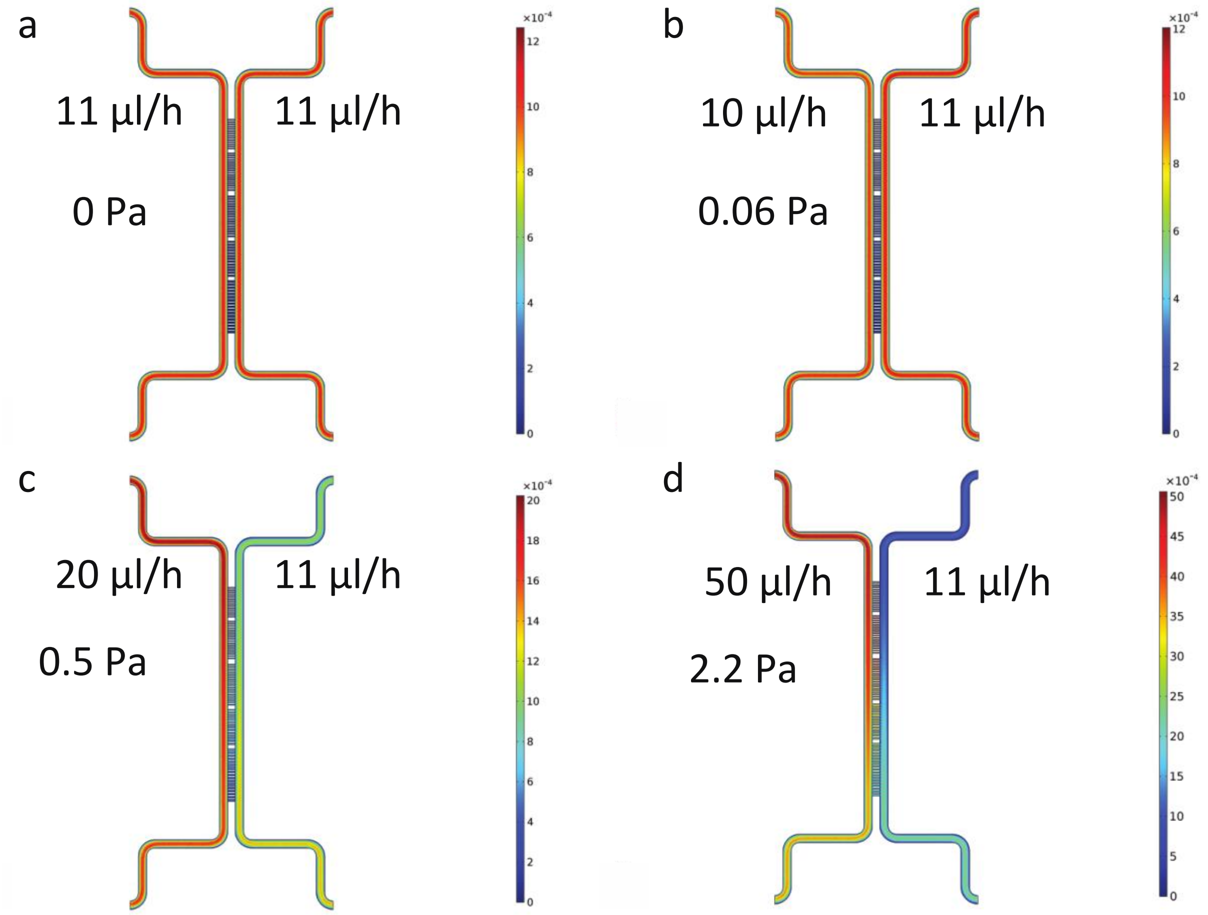


e

| !"#$ %&() *%µ"- . / | 01% & %00 | 31% & %00 | 41% & %00 |
|---|---|---|---|
| 5 67%8. ) ' &9' () %0- */ | ! " #$% | &$" #%% | ' ( ! " #%% |
| : 5 ; %8. ) ' &&() %0- */ | $$#) % | $! $#%% | &( ! &#%% |
| 5 67%8. ) ' &%(&) **%<' / | %#%) | %#! % | ' #( ) |
| : 5 ; %8. ) ' &%(&) **%<' / | %#( % | %#$) | &#( ! |

Figure S3: 3D simulation of the flow field at different flow conditions from COMSOL. The flow rates in the left and right chambers were, respectively, 11 – 11 (a), 10 – 11 (b), 20 – 11 (c), 50 – 11 (d) µl/h which gave an average shear stress of 0, 0.06, 0.5 and 2.2 Pa respectively within a narrow bridge as described in the main text.
